# Mean-field equations for neuronal networks with arbitrary degree distributions

**DOI:** 10.1101/118463

**Authors:** Duane Q. Nykamp, Daniel Friedman, Sammy Shaker, Maxwell Shinn, Michael Vella, Albert Compte, Alex Roxin

## Abstract

The emergent dynamics in networks of recurrently coupled spiking neurons depends on the interplay between single-cell dynamics and network topology. Most theoretical studies on network dynamics have assumed simple topologies, such as connections which are made randomly and independently with a fixed probability (Erdös-Rényi network) (ER), or all-to-all connected networks. However, recent findings from slice experiments suggest that the actual patterns of connectivity between cortical neurons are more structured than in the ER random network. Here we explore how introducing additional higher-order statistical structure into the connectivity can affect the dynamics in neuronal networks. Specifically, we consider networks in which the number of pre-synaptic and post-synaptic contacts for each neuron, the degrees, are drawn from a joint degree distribution. We derive mean-field equations for a single population of homogeneous neurons and for a network of excitatory and inhibitory neurons, where the neurons can have arbitrary degree distributions. Through analysis of the mean-field equations and simulation of networks of integrate-and-fire neurons, we show that such networks have potentially much richer dynamics than an equivalent ER network. Finally, we relate the degree distributions to so-called cortical motifs.

## INTRODUCTION

The dynamics in neuronal networks is strongly influenced by the patterns of synaptic connectivity. Networks in which connections are made randomly and independently with a fixed probability, so-called Erdös-Rényi (ER) networks, exhibit broad distributions of firing rates by virtue of the quenched variability in inputs coupled with the expansive nonlinearity of the neuronal transfer function in the fluctuation-driven regime [1, 2]. Allowing for clustered structure, in which neurons are more likely to be connected within clusters than across them, leads to multi-stable attractor states which may underlie working memory in cognitive tasks [3–5]. Analogously, networks in which the connectivity between neurons depends on the difference in their selectivity for a continuously varying quantity, e.g. orientation or spatial angle, can exhibit bump attractors [6, 7]. The detailed shape of such “spatially” structured connectivity (space may refer to feature space as opposed to distance along the cortical surface) in conjunction with synaptic delays determines the stability of such bumps and can lead to wave propagation [8].

Experimental data on patterns of synaptic connectivity in cortical slices indicates that cortical networks are not ER [9–11]. In particular, motifs between neuron pairs and triplets measured in experiment deviate from the expected values from ER networks, e.g. reciprocal connections are more frequent than expected. Additionally, it is found that neurons which share more common neighbors are more likely to be connected, a feature which has been attributed to the presence of clustering, e.g. [11], although this remains to be shown directly.

Inspired by these findings, recent theoretical work has looked at how changes in the frequency of second order motifs affects the synchronization of neuronal oscillators [12]. It has also been shown that allowing for broad degree distributions (distributions of the numbers of incoming and outgoing connections) can strongly affect dynamics in networks of asynchronous spiking neurons [13]. In that work, a heuristic firing rate equation was derived which showed that the in-degree distribution, which is related to the frequency of convergent connectivity motifs, strongly shapes the firing rate dynamics. On the other hand, the out-degree (divergent motifs) affects subthreshold correlations. However, the firing rate equations did not allow for correlations between in-degree and out-degree.

In this paper we derive and analyze heuristic mean-field equations for a network of neurons with arbitrary degree distributions (DDs), including those with correlated in-degree and out-degree. We first illustrate the derivation with a single population of neurons, and show that increasing the covariance between in-degree and out-degree is equivalent to increasing the strength of synaptic weights as far as the linear stability of fixed point solutions is concerned. We then derive the mean-field equations for a network of excitatory and inhibitory neurons (E-I network) and show that there are four relevant macroscopic variables: the four synaptic outputs *S_ee_*, *S_ei_, S_ie_*, S_*ii*_, as opposed to the two firing rates *r_e_*, *r_i_* traditionally used in mean-field descriptions of ER or all-to-all coupled networks. Finally, we show that the covariance between degrees is related to chain and reciprocal motifs, allowing one to use our model to study how such motifs shape the mean-field dynamics.

## A MEAN-FIELD MODEL FOR NETWORKS WITH ARBITRARY DEGREE DISTRIBUTIONS

We consider how network topology affects dynamics in large networks of recurrently coupled neurons. Specifically, we look at a measure of network connectivity called the degree distribution (DD), which is the distribution of numbers of pre-synaptic and post-synaptic inputs, or in-degree and out-degree of the cells, respectively. For simplicity we first derive a mean-field description for a single neuronal population of statistically homogeneous neurons. We then extend this framework for a canonical circuit which includes both excitatory and inhibitory neurons (E-I). Our main finding is that allowing for correlated DDs increases the dimensionality of the standard Wilson-Cowan firing rate description of an E-I circuit from two to four, increasing the range of possible network behaviors. This expansion in the underlying dimension occurs because the correct mean-field model for neuronal networks with correlated DDs is in terms of the synaptic output, of which there are four types (EE,EI,IE,II). Finally, we express the moments of the DDs in terms of so-called cortical motifs, which allows one to see at a glance the effect of certain motifs on mean-field dynamics in our framework.

### A mean-field description of a single neuronal population

In this section, we develop mean-field equations that describe the activity of a single statistically homogeneous population of neurons with degree distribution *ρ*(*x,y*), where *x* denotes (normalized) in-degree and *y* denotes (normalized) out-degree. We formulate a firing rate equation based on the Wilson-Cowan formalism by letting *r*(*x,t*) be the firing rate at time t of a neuron with indegree *x*. We start with the following heuristic equation rate equation (see *Appendix* for a derivation)

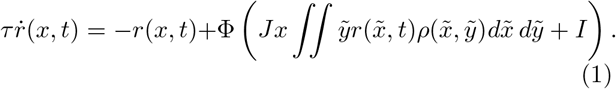

The key difference from typical Wilson-Cowan equations is that the sum over the network became the integral over the degree distribution *ρ*( *˜*, *˜*) with coupling strength *Jx ˜* that is proportional to the pre-synaptic neuron’s out-degree ˜ and the post-synaptic neuron’s in-degree *x*.

The firing rate equation, Eq.(1) is an infinite-dimensional system of equations, as it represents an equation for *r*(*x, t*) for all values of the in-degree *x*. Even if we transformed the equation to include just a finite number of in-degrees, using Eq. (1) directly would require keeping track of the firing rate for neurons of each in-degree. To more concisely capture the behavior of the population, we would like to develop a mean-field description of the population’s activity, i.e., develop a closed equation for a single population-averaged quantity.

A natural choice for a population-averaged quantity is the average of the firing rate over the population. We begin by attempting to derive an equation for the evolution of the average firing rate *R*(*t*) = 〈 *r*(*x,t*) 〉, where 〈•〉 denotes the average over the network: 〈*f*(*x,y*) 〉 = ∫*f*(*x,y*) *ρ*(*x,y*)*dx dy*. If we integrate each term in Eq. (1) over the distribution of degrees ρ(*x, y*), we obtain the following equation for the dynamics of the firing rate:

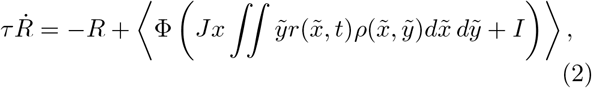

Unfortunately, the dependence of the recurrent input, or integral term, in Eq. (2) on *r*(*x, y*) is not in terms of the mean firing rate *R*(*t*). Knowing the value of the mean *R*(*t*) (and nothing more about the individual rates *r*(*x,t*)) is insufficient to determine the recurrent input, so we cannot close the system into an equation for just *R*(*t*).

There is one exception where we can develop a closed equation for the mean rate *R*(*t*): when the in-degree and out-degree are uncorrelated and hence the joint degree distribution factorizes into ρ (*x,y*) = ρ_*x*_(*x*) ρ_*y*_(*y*). In this case, the recurrent input depends on *r*(*x, t*) only through the mean *R*(*t*):

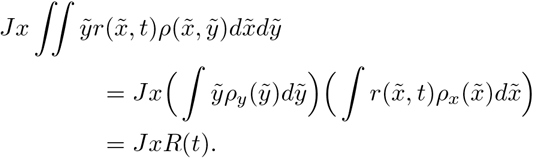

(Recall that *y* represents in-degree *normalized by the mean degree,* so 〈*y*〉 = 1.) In this special case, Eq. (2) closes to become a mean-field equation for *R*(*t*):

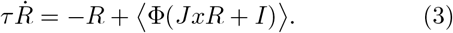

Eq. (3) and variants thereof for networks with delay and for E-I networks have been studied already in [13] [14]. The upshot is that changing the width of the in-degree distribution is akin to changing the gain of the fI curve Φ as far as linear stability is concerned. Therefore, networks with broad in-degree distributions can be more (or less) susceptible to instabilities, e.g. to oscillatory states. In this paper we go beyond this simple case and consider networks for which in-degree and out-degree may be correlated. If they are, then Eq. (3) is no longer valid.

In the general case with correlated degrees, we cannot express the mean firing rate self-consistently. In general, rather than being proportional to the mean rate, the recurrent input in Eq. (1) is proportional to the mean of the firing rates *r*(*x*˜, *t*) multiplied by the out-degree *y*˜.

Since, with correlated degrees, the out-degree y˜ depends on the in-degree *x˜* in *r*(*x*˜,*t*), we must change our perspective from averaging the firing rate alone to averaging the firing rate weighted by the number of outgoing connections. The form of Eq. (1) dictates that the relevant mean-field variable is actually the firing rate weighted by the number of outgoing connections

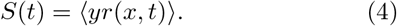

We will call *S*(*t*) the *synaptic drive* as it represents the average activity of the synapses, or the edges in the network. It's the synaptic drive *S*(*t*), rather than the mean firing rate *R*(*t*), that is the correct projection of the firing rates *r*(*x, t*) in the presence of correlated degrees, allowing us to derive a closed mean-field equation.

If we average each term in Eq. (1), weighted by the out-degree, we obtain a self-consistent equation for the synaptic drive

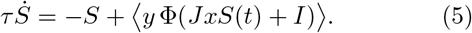

This mean-field synaptic drive equation for the single quantity *S*(*t*) captures the influence of an arbitrary degree distribution *ρ*(*x, y*) on the dynamics of the network. (Recall that the mean 〈•〉 is an average over *ρ*(*x,y*).) Once the synaptic drive *S*(*t*) is determined from Eq. (5), one could reconstruct the mean firing rate *R*(*t*) from Eq. (2) or even the distribution of firing rates *r*(*x, t*) as a function of in-degree from Eq. (1). Both these equations are driven solely by the synaptic drive, as they can be rewritten as

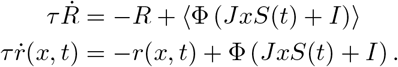

If the fI curve Φ were linear, Eq. (5) could be greatly simplified by noting that

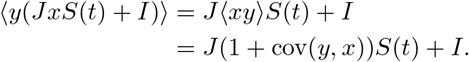

This calculation reveals that the covariance of the indegree and out-degree modulates the strength of the recurrent input, creating an effective coupling strength of *J*(1 + cov(*y,x*)). (Note that this covariance is normalized by the square of the mean degree *d*^2^; the effect of the mean degree is already included in *J*.) As a consequence, a positive covariance (indicating that the neurons that receive many inputs also have many outputs) is equivalent to having stronger synapses, as far as the first order effect on mean rates goes. A strong negative covariance (indicating that those neurons with many outputs receive but few inputs) diminishes the recurrent input and, in an extreme case, could essentially eliminate the effects of the coupling. (Since *x* and *y* are non-negative with mean 1, we can conclude that cov(*y, x*) ≥−1).

For nonlinear fI curves, the effect of the degree covariance can no longer strictly be reduced to multiplying the effective coupling strength by 1 + cov(*y,x*). However, a similar effect of the covariance on the stability of steady state solutions can be seen. We consider the stability of a steady-state synaptic drive *S*_0_ with the ansatz *S*(*t*) = *S*_0_ + *δSe^λt^*, where *δS* ≪ 1. Plugging this expression into Eq. (5) yields the characteristic equation for the eigenvalue *λ*

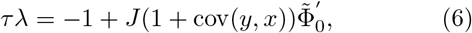

where

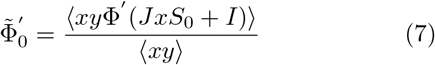

is an “effective gain”, a weighted average of the gains Φ ′. This result is comparable in form to the characteristic equation of a standard one-population Wilson-Cowan equation with equivalent notation, which is *τλ* = −1 + *J*Φ′_0_, where Φ′_0_ is the gain at the steady state. For the standard Wilson-Cowan equation, an instability occurs if the product *J*Φ′_0_ of the synaptic weight and the gain of the nonlinear transfer function at the steady state is greater than one. The characteristic equation for arbitrary degree distributions, Eq. (6), reflects two new features introduced by the network structure. The first is the introduction of the “effective gain” Φ̃′ of Eq. (7). Ref. [13] studied how the degree distribution shapes this gain when the degrees were uncorrelated; correlations between degrees may introduce additional effects. The second new feature is the dependence of the stability on the covariance of the degrees through the modification of the effective coupling strength to *J*(1 + cov(*y, x*)), as described in the case with a linear fI curve.

### Mean-field equations for an EI network

We extend the mean-field analysis to a network of two populations straightforwardly. Since the EI firing rate equations of Eqs. (27) are in the same form as the single population firing rate equations of Eq. (26), we would run into the same difficulty if we attempted to close the equations in terms of the two population average firing rates. Instead, the equations dictate that the correct projection of the firing rates is again in terms of synaptic drives. With two populations, however, we obtain four different synaptic drive variables.

The synaptic drive variables correspond to the four recurrent input integrals of Eqs. (27). Let 〈•〉_*a*_ denote an average over the degree distribution *ρ_a_* (**x**_*a*_, **y**_*a*_) of population *a*:

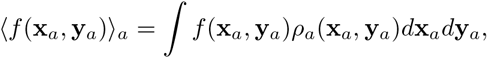

where **x**_*a*_ = (*x_ae_, x_ai_*) denotes the E in-degree and I indegree to a neuron in population a and **y** = (*y_ea_,y_ia_*) denotes the E out-degree and I out-degree from a neuron in population *a*. (See the appendix for more details on the two population degree distribution.) Then, the synaptic drive from population b onto population *a* is the firing rate of population *b* weighted by the corresponding out-degree:

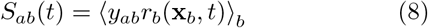

With correlated degree distributions, we require four equations, rather than the typical two, to capture the dynamics of two population. This requirement stems from the fact that, with correlated degree distributions, the dynamics of each type of input can be different for the excitatory and inhibitory populations. For example, the excitatory synaptic drives *S_ee_* and *S_ie_* to the excitatory and inhibitory population would have identical dynamics only for special network structures for which the integrals of Eq. (9a) and Eq. (9b) were identical, i.e., the integrals were invariant to the distinction between the E out-degree *y_ee_* and the I out-degree *y_ie_* of the excitatory population. Below, we show that differences among the correlations between in- and out-degrees are sufficient to create four independent equations.

### Linear Stability

We would like to know if the mean-field Eqs. (9) exhibit richer dynamics than the standard two-dimensional firing rate equations commonly used to describe the activity in E-I networks. As a first step we consider the linear stability of fixed point solutions. As in the case of a single population, linearizing about the fixed-point solution leads to terms which are proportional to the product of an effective gain and a covariance of degrees. The difference with the simple, one-population model is that in the two-population model now there are eight covariance terms; see Fig. 1.

**Fig. 1.**
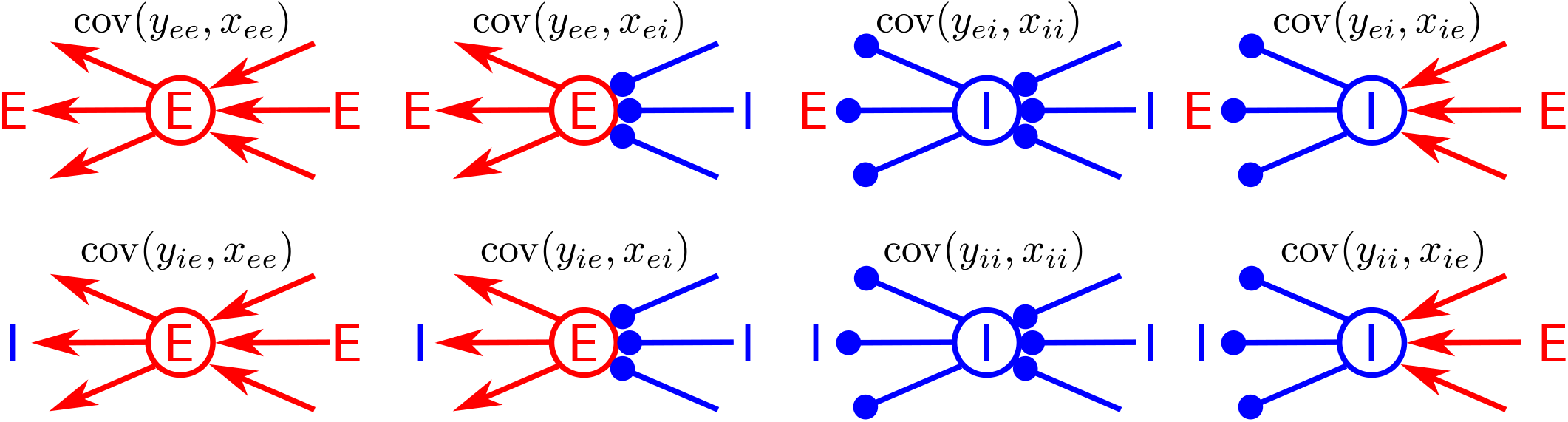
The eight degree covariances in a network of excitatory and inhibitory neurons. In our analysis, we include the effect of the covariance cov(y_ab_,x_bc_) in the parameter β_abc_; see Eq. (13).

Each of the four equations in Eqs. (9) depends on two of those covariances, as determined by the *x_ab_* and *y_ab_* factors in the equation. In general, these eight covariances in the network structure will differ from one another. As a result, each of the four synaptic drives will have different dynamics and the dynamics will be truly four-dimensional.

For example, consider the excitatory synaptic drives These four scalar quantities for *a, b* ∈ {*e, i*} capture the averages of the firing rates that determine the dynamics of the network. We can write self-consistent equations for these drives by multiplying the equations from Eq. (27) by the appropriate out-degree and integrating:

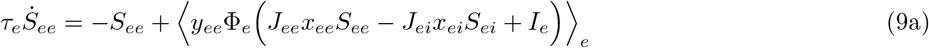

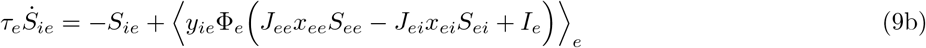

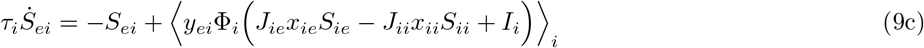

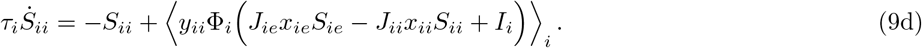

*S_ee_* and *S_ie_*. In the standard firing rate equations, there would be only one excitatory variable, the average firing rate of the excitatory population. Depending on the network structure, as captured by the covariances of Fig. 1, the excitatory synaptic drive could be different for the excitatory population (*S_ee_*) than for the inhibitory population (*S_ie_*). The first column in Fig. 1 illustrates one pair of covariances that could lead to a difference in those excitatory synaptic drives. The top of the first column illustrates the covariance between the E in-degree and the E out-degree of excitatory cells (cov(*y_ee_*,*x_ee_*)). On the other hand, the bottom of the first column illustrates the covariance between the E in-degree and the I out-degree of excitatory cells (cov(*y_ie_*, *x_ee_*)).

Imagine a network where these two covariances were significantly different, say where cov(*y_ee_, x_ee_*) > 0 and cov(*y_ie_, x_ee_*) < 0. In this case, those excitatory neurons which receive the most recurrent excitatory input would (A) project broadly to other excitatory cells and (B) project little to inhibitory cells. On the other hand, those excitatory neurons which receive the least recurrent excitatory input would (A) project little to other excitatory cells and (B) project broadly to inhibitory cells. As a result, the excitatory and inhibitory populations sample the excitatory synaptic output in very different ways. This effect is not equivalent to any change in the average synaptic weights *J_ee_* and *J_ie_*; it requires correlations among the network edges as captured by the covariances between in- and out-degrees (or similar correlations among the synaptic weights between individual neurons in a weighted network).

As with the single population case, the covariances hidden in the equations are revealed when one linearizes Eq. (9) to determine the stability of a steady-state synaptic drive 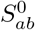. Since each equation is of the form,

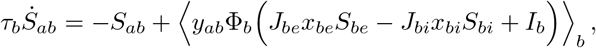

we can replace 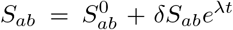and linearize the integral term as

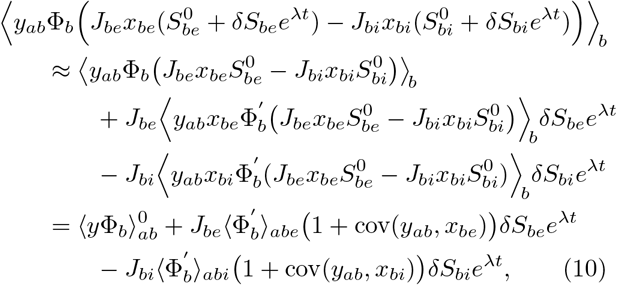

where 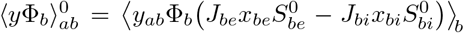 is the value of this integral at the steady state and

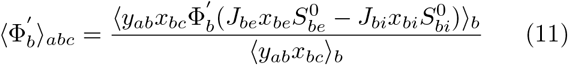

is the effective gain of population b, averaged and normalized with respect to the covariance of the in-degree from population c and the out-degree to population *a* for *a, b*, *c ∈* {*e,i*}. Hence, the triple subscript *eei,* for examples, represents the covariance between the I in-degree and E out-degree of an excitatory neuron, a covariance of degrees that influences the number of chains of connections from an inhibitory neuron onto excitatory neuron onto another excitatory neuron. For brevity in notation, we will combine the effective gain and covariance terms into a single parameter,

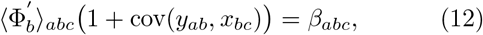

which is the effective gain of population *b*, combined with the covariance of its in-degree from population *c* and out-degree to population *a.* Note that *β_abc_* ≥ 0.

We will consider how changes in this parameter can affect the linear stability of fixed point solutions. In doing so, we will focus exclusively on the role of changes in the covariance, that is, we will consider the effective gain to be fixed. This assumes, for example, that as the covariances in the degrees are varied additional parameters must be adjusted (such as the external input) to maintain constant gain. This is a reasonable assumption given that what is observable and hence known in cortical circuits is the pattern of activity, e.g. mean firing rates, and not the patterns of connectivity. Therefore we will be investigating the effect of changes to the connectivity, while holding the level of activity, and hence gain, fixed.

Using this notation, the linear stability analysis leads to the following characteristic equation for the eigenvalue *λ*:

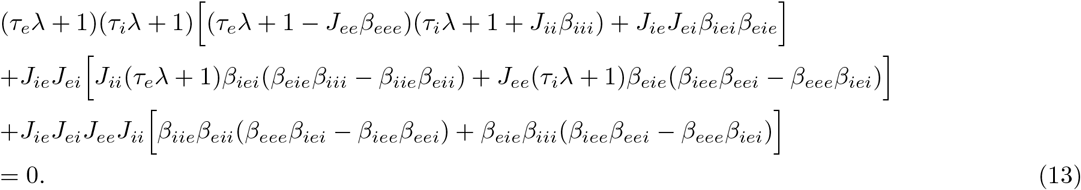

To compare to typical characteristic equations of Wilson-Cowan models, imagine a special case of the network structure where the integrals Eq. (9a) and Eq. (9b) were identical and the integrals of Eq. (9c) and Eq. (9d) were identical. In that case the excitatory synaptic drives would be identical, *S_ee_* = *S_ie_* and the inhibitory synaptic drives would be identical, *S_ei_* = *S_ii_,* reducing Eqs. (9) to two equations. In order for these equations to be identical, the top row of covariances in Fig. 1 must be identical to the bottom row, i.e., β_*eab*_ = β_*iab*_ for any pair of populations *a* and *b.* For this special case, we can drop the first index of β and just refer to it as *β_ab_.* With this simplification, the combinations of β’s in parentheses cancel due to symmetry, and the characteristic equation Eq. (13) simplifies to just its first term

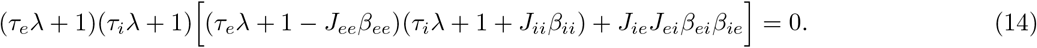

The factor in the square brackets is essentially the characteristic equation for a standard Wilson-Cowan description for an E-I network.

In the full characteristic equation, Eq. (13), the second and third terms reflect effects from having the covariances of first row in Fig. 1 differ from those of the second row, i.e., they capture the effects of having the inhibitory and excitatory populations sample the recurrent excitation and inhibition in different ways, as quantified by different degree covariances. We will provide an example of such an effect in the next section, without doing an exhaustive analysis.

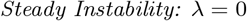

A steady instability indicates a saddle node bifurcation and a region of bistability, which can be computationally relevant for modeling working memory states. For E-I networks that reduce to two-dimensional dynamics (which include standard Wilson-Cowan E-I networks), setting λ = 0 in Eq. (14) indicates that a steady instability occurs only when the recurrent excitatory input is sufficiently strong, i.e. when *J_ee_ β_ee_* exceeds a critical value. For standard Wilson-Cowan E-I networks, this instability can occur either by increasing the strength *J_ee_* of the recurrent excitatory synaptic weights, or increasing the gain, for example through external inputs (as *β_ee_* includes only the gain factor when one ignores degree covariances). Without recurrent excitation it is not possible to have steady instabilities and hence working memory states in standard Wilson-Cowan equations for E-I networks. The same conclusion is true even for networks with strong degree correlations, as long as the dynamics remain two-dimensional due to the top row of covariances in Fig. 1 being identical to the bottom row.

In general, however, the presence of correlated DDs in E-I networks can actually allow for working memory states even in the absence of recurrent excitation. To see this, we set *J_ee_* = 0 in Eq. (13), which (for *λ* = 0) gives

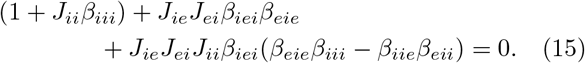

Since the parameters *J* and *β* are non-negative, we must have *β_iie_β_eii_* > *β_eie_β_iii_* in order for there to be a steady instability. What does this condition mean? To begin, notice that *β*_*eie*_ and *β*_*iie*_ represent the top and bottom covariances of the fourth column of Fig. 1, respectively; *β*_*eii*_ and *β*_*iii*_ represent the top and bottom covariances of the third column of Fig. 1, respectively. We have already determined that we need the covariances in the top and bottom rows of Fig. 1 to differ in order to obtain bistability in the absence of recurrent excitation. The necessary condition for bistability, *β_iie_β_eii_* > *β_eie_β_iii_,* specifies in what manner these covariances must differ for the bistability.

To begin, let’s focus on the covariances from the fourth column of Fig. 1. If we have a network where *β_iie_* > *β_eie_* and the covariances in the third column are equal, the necessary condition for bistability is met. This pattern of covariances indicates that inhibition onto inhibitory neurons is stronger than the inhibition onto excitatory neurons, again not because of different synaptic weights, but rather because E and I are sampling inhibitory outputs in very different ways. When *β_eie_* is small, E cells would tend to receive inhibition from I cells which receive little excitation. At the same time, when *β_iie_* is large, I cells would receive inhibition from strongly excited I cells. This pattern is illustrated in the right column of Fig. 2.

**Fig. 2.**
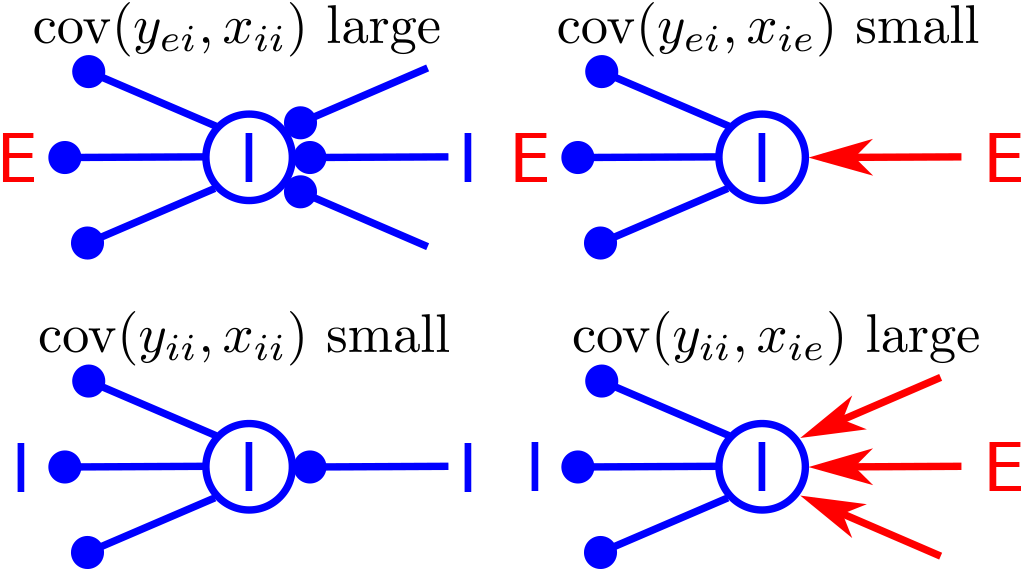
A situation conducive to working memory states, even in the absence of recurrent excitation. The top row illustrates that E cells tend to receive inhibition from neurons which are weakly excited and strongly inhibited. On the other hand, as illustrated by the bottom row, I cells tend to receive inhibition from neurons which are strongly excited and weakly inhibited. This situation is consistent with the condition *β_iie_*, *β_eii_* > *β_eie_ β_iii_*, which is necessary to satisfy Eq. (15).

An alternative pattern to meet the necessary condition for bistability involves the covariances from the third column of Fig. 1: a network where *β_eii_* > *β_iii_* while the covariances in the fourth column are equal. This pattern would mean that E cells tend to receive inhibition from I cells which are strongly inhibited, with the opposite pattern for I cells, as illustrated in the left column of Fig. 2.

This scenario of working memory states through dis-inhibition of excitatory neurons has been explored before when there are two distinct inhibitory populations, e.g. [15]. Here we have a similar mechanism, which however relies on distinct ways of sampling a single highly heterogeneous population of inhibitory neurons.

### Simplified mean-field model

The mean-field equations Eqs. (9) are unwieldy due to the integration over the joint degree distributions represented by the 〈•〉_*a*_ The integration essentially amounts to a re-scaling and translation of the nonlinear transfer function Φ. We can write down simplified mean-field equations with an effective nonlinear transfer function with the same overall structure as Eqs. (9).

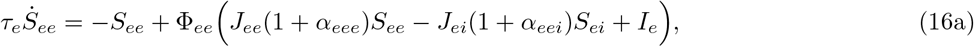

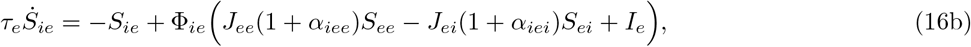

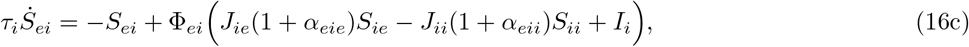

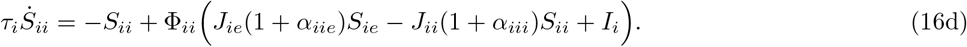

Specifically, we choose the nonlinear transfer functions Φ so that Eqs. (16) have an identical fixed point structure and linear stability as in Eqs. (9). Furthermore, as long as the DD is unimodal the effective nonlinearity in Eqs. (9) is monotonically increasing and hence qualitatively similar to that in Eqs. (16). The resulting equations are in the same form as standard Wilson-Cowan equations for four populations, two excitatory and two inhibitory, with a particular form of coupling given by the parameters *J_ab_*(1 + *α_cab_*). The parameters *α* are chosen so that the linear stability of perturbations of fixed points in Eqs. (16) is given by Eq. (13) with *β_cab_* = Φ′_*ca*_ (1 + *α_cab_*). Given the definition of β in Eq. (12), we see that parameters are essentially

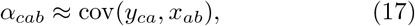

i.e., α_*cab*_ represents the covariance between the in-degree *x_ab_* and the out-degree *y_ca_*.

One advantage of the simplified equations is that their bifurcation diagrams can be quickly computed once one chooses a form of the nonlinearities Φ. Fig. 3 shows an example of a bifurcation diagram for Eqs. (16) when *J_ee_* = 0, in which case *S_ee_* can be eliminated and the system reduces to three dimensions. (See the figure caption for parameter values.) When all of the degree covariances are the zero, i.e. α_*cab*_ = 0 for all combinations of *a*, *b* and *c*, then Eqs. (16) actually reduce to only two equations and bistability is not possible without recurrent excitation, as seen earlier. Bistability occurs when the covariance between the E in-degree and I out-degree of inhibitory neurons is increased so that *α_iie_* = 1. This parameter choice corresponds to increasing the covariance in the lower right of Fig. 2, which is conducive to bistability.

**Fig. 3.**
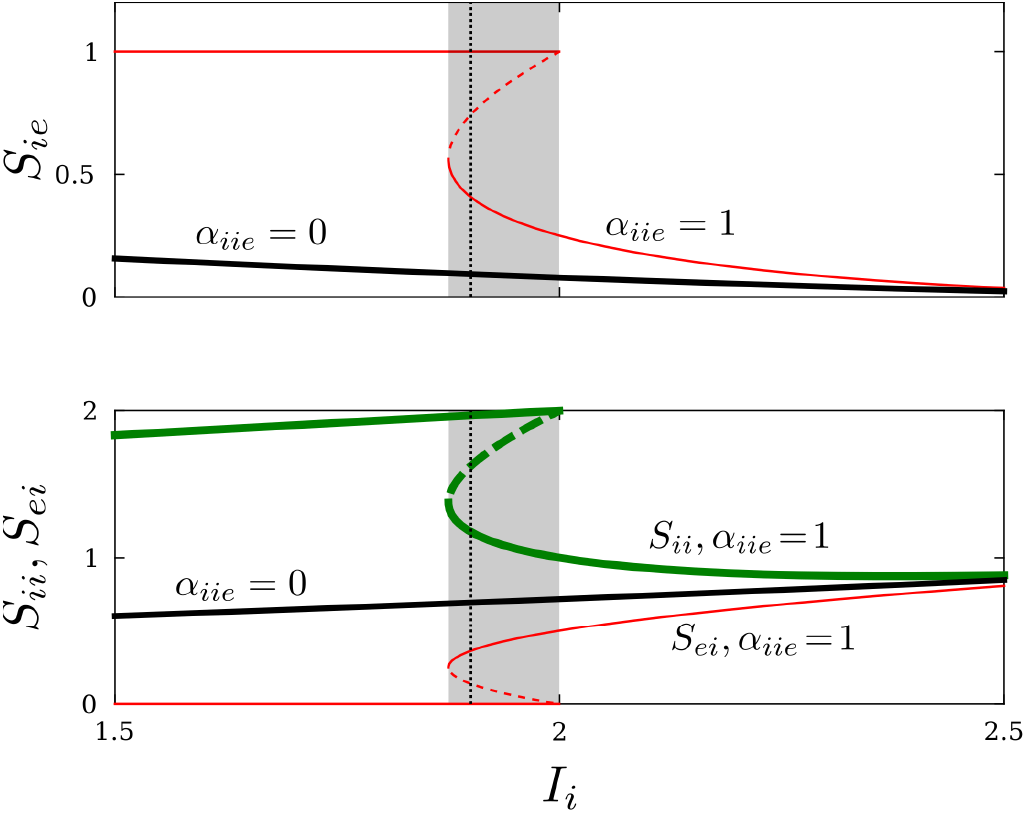
An example of bistability without recurrent excitation. Shown is a bifurcation diagram of fixed point solutions to Eqs. (16) for both a case without bistability (black lines) and a case with bistability (colored lines). Solid lines correspond to stable equilibria and dashed lines to unstable equilibria. Top: Black line *S_ie_* for α_*iie*_ = 0. Thin red line S_ie_ for *aue* = 1. Bottom: Black line *S_ei_* and *S_ii_* for α_*iie*_ = 0. (These synaptic drives are indistinguishable in this case.) Thin red (thick green) lines: *S_ei_* (*S_ii_*) for α_*iie*_ = 1. The dotted vertical line indicates the value of *I_i_* for which the simulations in Fig. 4 were done. The bistable region is shaded. Parameters: *α_abc_* = 0 for all *a, b, c* ∈ {*e, i*} except where noted, *I_e_* = 1, *τ_e_* = *τ_i_* = 1, *J_ee_* = 0, *J_ei_* = 1, *J_ie_* = *J_ii_* = 2, Φ_*ee*_(*x*) = Φ_*ie*_(*x*) = 0, *x*^2^ or 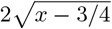 if *x* ≤ 0, 0 < *x* < 1 and *x* ≥ 1 respectively. Φ_*ei*_ (*x*) = Φ_*ii*_ (*x*) = 0 if *x* < 0 and *x* otherwise.

A sample simulation of bistable behavior is shown in Fig. 4. At time *t* = 200 the external input to the excitatory neurons is increased transiently, causing an increase in excitatory output *S_ie_* (top panel). There is an accompanying drop in the inhibitory output to excitatory cells *S_ei_*, although the synaptic inhibition onto inhibitory cells increases *S_ii_.* Finally, an increase in the external input to inhibitory cells at *t* = 400 causes *S_ie_* to drop back down to a low-activity fixed point.

**Fig. 4.**
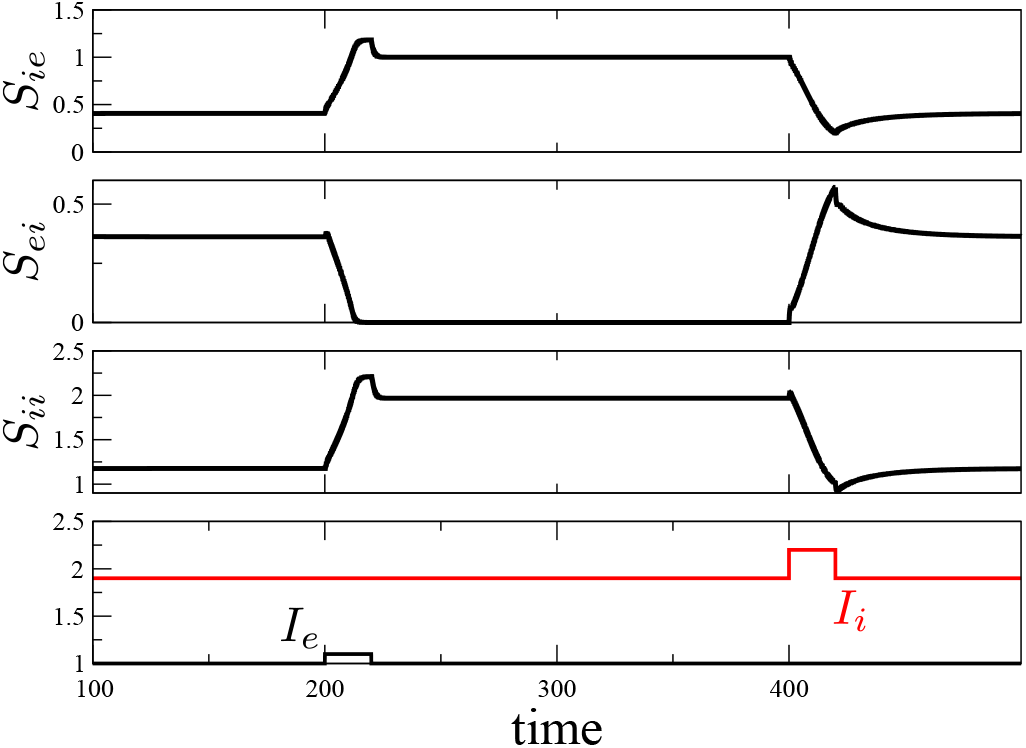
A sample simulation in the bistable regime. The value of *I_i_* = 1.9 (see vertical dotted line in Fig. 3).

### Integrate-and-fire simulations

To further demonstrate how degree correlations can bestow bistability even without recurrent excitation, we simulated a network of 4000 excitatory and 1000 inhibitory integrate-and-fire neurons without E to E connectivity, see *Appendix* for details of the network and simulations. We coupled the neurons using a SONET network, a network model that allows one to generate networks with prescribed degree correlations. In the SONET model, one specifies the number of different types of chains, which corresponds to the degree covariances, as shown below. In particular, the SONET chain parameters 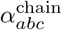 correspond to the *α_abc_* in the simplified mean-field model above.

If we set all chain parameters 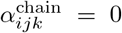, the network does not display bistability, consistent with the theory. To generate a network with appropriate covariances for working memory, we used parameters based on the right column of Fig. 2, which lead to the degree distribution of the inhibitory population shown in Fig. 5. We reduced the number of E to I to E chains, setting 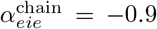, yielding degree correlations depicted in the upper right of Fig. 2 and evident in the upper right of Fig. 5. We increased the number of E to I to I chains, setting 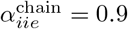, yielding degree correlations depicted in the lower right of Fig. 2 and evident in the lower right of Fig. 5.

**Fig. 5.**
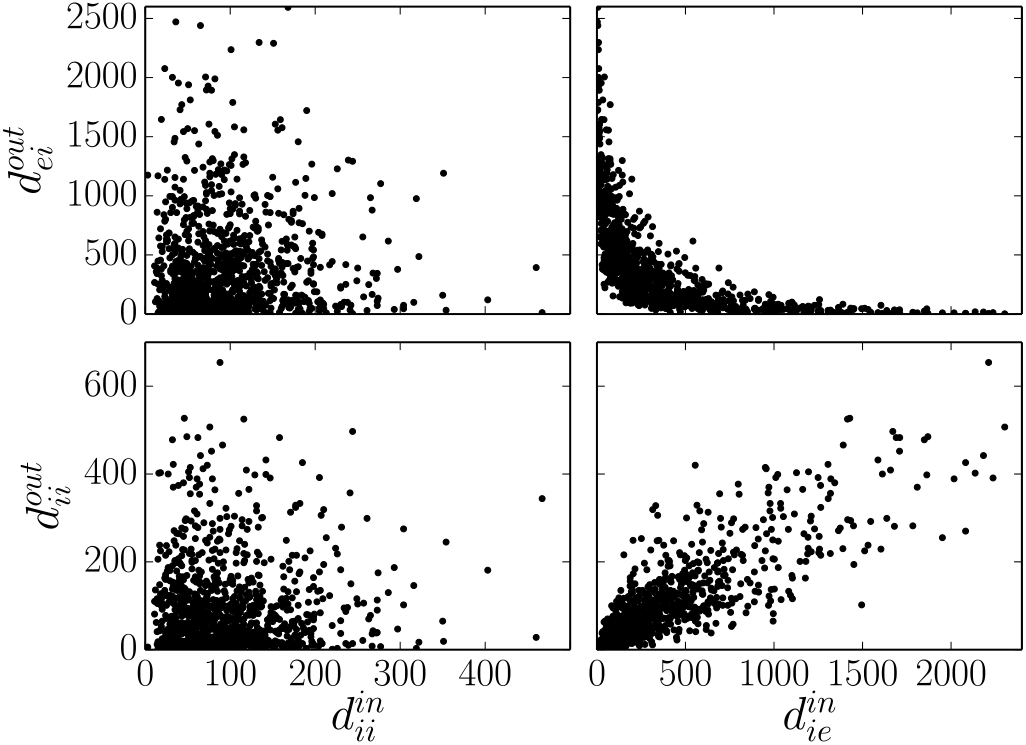
The inhibitory population degree distribution of the network underlying bistability induced by degree correlations. Since 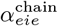 is small the E out-degree and E in-degree of the inhibitory population are highly anti-correlated (lower right, c.f., lower right of Fig. 2). Since 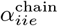 is large, the I out-degree and E in-degree of the inhibitory population are highly correlated (upper right, c.f., upper right of Fig 2). The I indegree is uncorrelated with the out-degrees of the inhibitory population (left column).

Fig. 6 demonstrates the presence of bistability in this network analogous to Fig. 4. The excitatory population can be switched from a high firing mode to a low firing mode with transient changes to its input rate. Fig. 7 illustrates that this bistability is facilitated by a shift of the firing pattern of inhibitory neurons. During the high firing mode, the only inhibitory neurons that fire rapidly are those that receive a large amount of excitation and project minimally to the excitatory population. As shown in the lower right of Fig. 5, these neurons also tend to project broadly to other inhibitory neurons. Hence, in the high firing rate mode, inhibitory neurons that project broadly to excitatory neurons are silenced, allowing the excitatory population to fire more quickly.

**Fig. 6.**
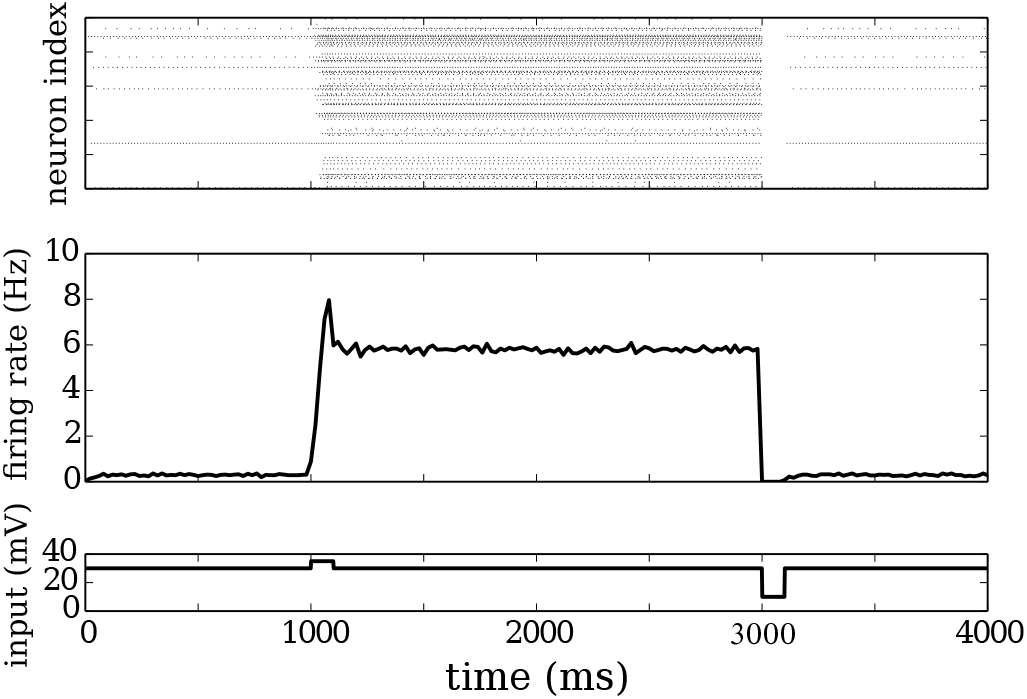
Integrate-and-fire network without recurrent excitatory coupling demonstrating bistability due to degree correlations. Top: raster plot of 50 neurons from the excitatory population. Middle: average firing rate of excitatory population. Bottom: input to excitatory population *RI_e_.* Integrate-and-fire parameters: *E_r_* = –65 mV, *E_e_* = 0 mV, *E_i_* = –70 mV, υ _th_ = –50 mv, υ_reset_ = –60 mV, τ _*e*_ = 20 ms, *τ_i_* = 10 ms, *T τ_a_* = 2 ms, *τ_g_* = 10 ms, *τ*_ref_ = 1 ms, *J_ei_* = 0.2, *J_ie_* = 0.1, *J_ii_* = 0.013, and *RI_i_* = 25 mV. Network parameters: *p_ee_* = 0, *p_ei_* = *p_ie_* = *p_ii_* = 0.1, 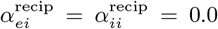, 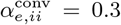, 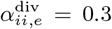, 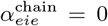, 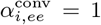, 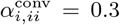, 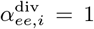, 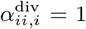, 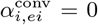, 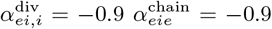, 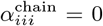, 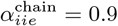, 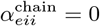.

**Fig. 7.**
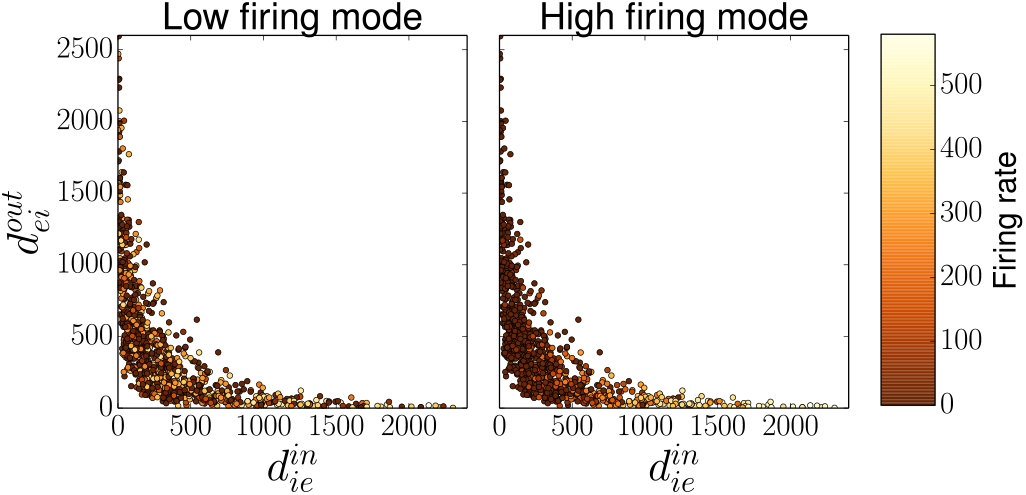
Average firing rates of inhibitory neurons illustrated as a function of their degree. Left: During the low firing rate mode (first or last second of simulation), the firing rates of individual inhibitory neurons are not strongly correlated with their degree. Right: During the high firing rate mode (middle period of simulation), only the inhibitory neurons with a high E in-degree and a low E out-degree are firing rapidly.

### Degree covariances are proportional to the number of chains in the network

We have seen that the dynamics in E-I networks can be strongly shaped by the presence of correlations between DDs. What do these correlations mean in terms of simple connectivity motifs, which can be determined straightforwardly in slice experiments? In fact, positive covariances indicate the presence of added chain motifs, above and beyond what is expected in an Erdo ös-Re ényi network. In the SONET model, we manipulated the number of chain motifs to create degree correlations. In general, one can show that a network with positive degree correlations has over-represented chain motifs.

This relationship between chains and degree correlations is illustrated in Fig. 8 which shows a chain motif from neuron *i* to k through *j*. We can calculate the probability of finding this particular motif based on the degrees of the neurons. In particular, the probability that cell *i* connects to cell *j* is just proportional to the product of the degrees 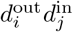, while the probability that *j* connects to k is proportional to 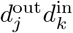. The probability of this chain motif in a network of *N* ≫ 1 neurons is then

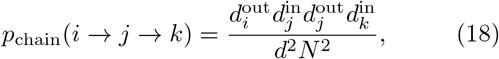

where *d* is the mean degree. Assuming independence between degrees of neighboring nodes (i.e., neglecting any assortativity or other higher order correlations), the expected value of this probability in the network is

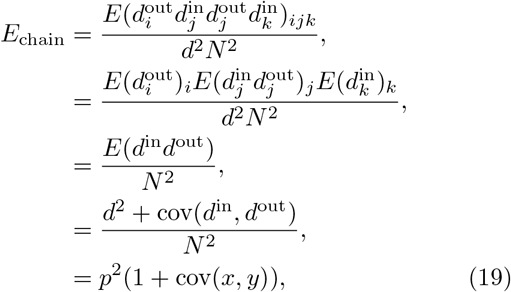

where *p* = *d/N* and (*x, y*) are the degrees normalized by the mean degree *d*. When the covariance is zero, then the probability of a chain motif is just *p*^2^, whereas increasing or decreasing covariance increases and decreases the probability of chains respectively. Therefore we can study the affect of different chain motifs on the mean-field by directly varying the covariances. In an E-I network, the number of chains from cell type *A* → *B* → *C* where *A,B,C* ∈ {*E, I*} is varied by changing the covariance cov(*y_CB_*, *x^BA^*).

**Fig. 8.**
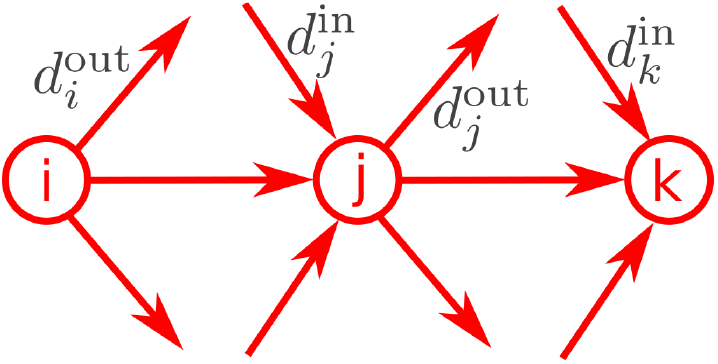
The probability of a chain motifs depends on the degrees. In this example there is a chain from excitatory neuron *i* to *j* to *k*. This particular motif depends on the four degrees 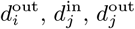 and 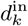.

Finally, we remark that reciprocal motifs are also a type of chain which goes from *i* → *j* → *i*. A calculation analogous to Eq. (19) shows that the probability of reciprocal connections also depends on the covariance as *E*_recip_ = *p*^2^(1 + cov(*x,y*))^2^. We also note that the probabilities of convergent and divergent motifs are proportional to the variance of the in-degree and the out-degree respectively [12, 13]. Therefore, a large fraction of chain motifs in such networks necessarily implies large fractions of convergent and divergent motifs as well.

## DISCUSSION

In this paper we have derived mean-field equations for a network of neurons with a given degree distribution. The formulation is based on the simple idea that the probability of a connection between two cells should be proportional to both the out-degree of the pre-synaptic cell and the in-degree of the post-synaptic cell. Indeed, in a network where connections are made randomly and with equal probability conditioned on degree, the probability that a single given out-going synapse from a cell *j* is made onto a cell *i* must be proportional to 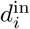, the in-degree of cell *i*. To obtain total probability of a connection from *j* to *i*, we multiply by 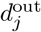, the number of out-going synapses from cell *j*. The overall connection probability is hence proportional to the product of degrees, 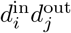. This approximation is reasonable as long as the number of neurons is large compared to the mean degree in the network.

One key consequence of this work is the discovery that the resulting relevant mean-field variables are not the firing rates of the neurons, but rather their synaptic output. As the number of different types of synaptic connections scales with the square of the number of populations, the effect of network structure is to dramatically increase the dimension of the mean-field dynamics.

For example, in a canonical E-I network, with two populations of neurons, incorporating the effects of the network structure leads to four-dimensional mean-field equations, in contrast to the standard two-dimensional mean-field equations of an Erdo ös-Re ényi random E-I network. Fig. 9 illustrates the interactions among the four mean-field variables *S_ee_*, *S_ie_*, *S_ei_* and *S_ii_*, which we call synaptic drives. These synaptic drives represent the edges of the original network, but they become the nodes of the effective graph (Fig. 9, right). Using the simplified description of Eq. (16), each edge in the effective graph corresponds to one of the eight different types of covariance α_*cab*_ (Eq. (17)) of the degree-distribution. Since these degree-covariances correspond to chains of two edges in the original network, we see that these two-edge chains become the edges of the effective graph describing the network dynamics.

**Fig. 9.**
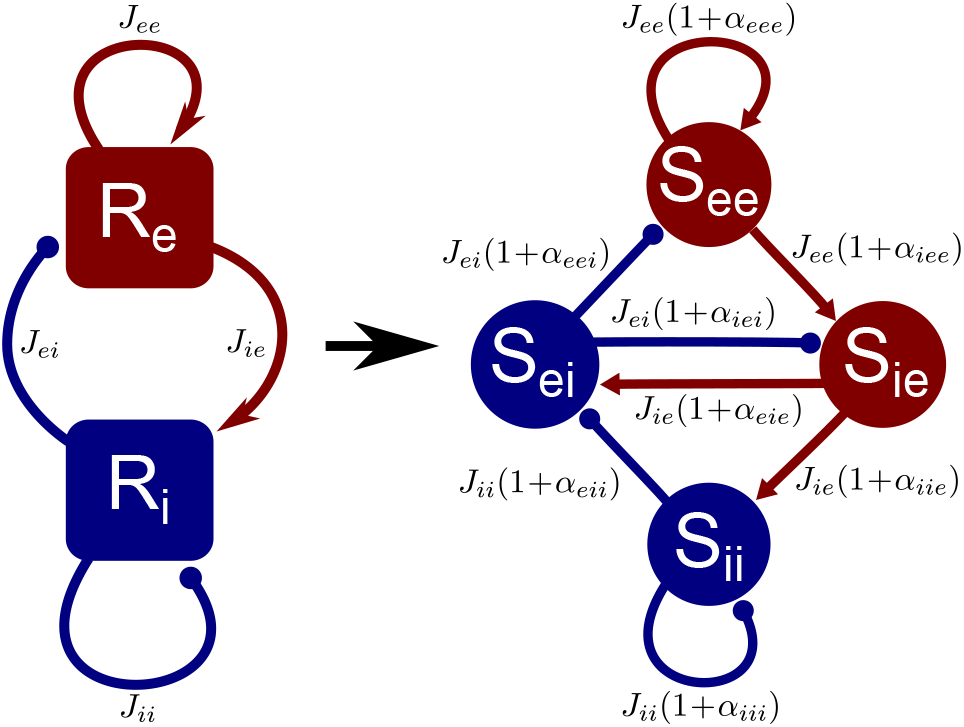
Illustration of the effective mean-field network obtained from an EI network with arbitrary degree distributions. Left: for an Erdo ös-Re ényi random network, one can obtain two-dimensional mean-field dynamics of the firing rates *R_e_* and *R_i_* of the excitatory and inhibitory population, coupled by an average connectivity *J_ab_*. Right: in general, the presence of correlations between neurons’ in- and out-degrees creates four-dimensional mean-field dynamics. One can represent these equations as a network with the four synaptic drives *S_ee_*, *S_ie_*, *S_ei_* and *S_ii_* as the nodes. The coupling among these nodes is determined not just by the average connectivity *J_ab_*, but also the correlations α_*abc*_ among the in-and out-degrees, as captured in Eq. (16).

These results indicate that given certain types of degree distributions, the excitatory (or inhibitory) synaptic output to excitatory neurons can be radically different from the output to inhibitory neurons. Fig. 9 illustrates how this effect is not equivalent to changing average synaptic weights. For example, in the standard E-I firing rate model (Fig. 9, left), when the synaptic weights *J_ii_* and *J_ei_* are different, then I cells and E cells receive different amounts of synaptic inhibition, in *magnitude.* However, the time course of the synaptic input is identical for both populations, as it simply given by the time course of the single dynamical variable *R_i_*. In the case of our mean-field equations (Fig. 9, right), the synaptic outputs *S_ii_* and *S_ei_* are *dynamically* independent variables.

It may seem counterintuitive that we obtain more dynamic variables than classes of neurons in the original network. However, the dynamics of different pooled outputs of a single population can have distinctly different dynamics if that population is highly heterogeneous. For example, the I cells illustrated in Fig. 7 have highly heterogenous behavior, depending on their degrees. Since E cells and I cells received their input primarily from different “sub-classes” of I cells, these two inhibitory inputs were dynamically independent. In reality, there are no sub-classes in our model; merely, the broad continuum of degrees leads to some I cells that receive many E inputs and some I cells that receive very few E inputs. For this example network, E cells received their inhibition from I cells that received few E inputs, while I cells received their inhibition from I cells that received many E inputs, i.e., the two population sampled the activity of very different I cells. This scenario can be summarized succinctly by stating that the covariance cov(*y_ei_,x_ie_*) is small and cov(*y_jj_, x_ie_*) is large, which is in fact conducive to working memory-like bistability in E-I circuits. (See the right-hand column of Fig. 2.)

In fact, in our network any such “sampling-of-different-sub-classes” argument relies on there being a significant difference in covariances related to differential inputs between E and I cells. In a classic E-I Erdoös-Reényi random network all covariances are zero and a description of the mean-field activity in terms of firing rates alone is possible. Similarly, in a network in which DDs are broadly distributed, if in- and out-degrees are independent then a mean-field description in terms of firing rates is self-consistent [13]. Even in the presence of significant covariances between degrees, the mean-field dynamics will be two-dimensional as long as the degree distribution has symmetries that ensure the synaptic output does not depend on the target population. At the level of the simplified dynamics captured by Eqs. (16) and 17, the symmetries are the requirement that degree covariances do not depend on the type of output from the neuron, i.e., cov(*y_eb_, x_bc_*) = cov(*y_ib_, x_bc_*) for all b, c, ∈{*e, i*}this case, Eqs. (16) would simplify to two equations for *S_e_* = *S_ee_* = *S_ie_* and *S_i_* = *S_ei_* = *S_ii_;* the four-node effective graph of the right of Fig. 9 would collapse to a two-node graph in terms of *S_e_* and *S_i_*.

Recent work has studied the spectrum of random matrices for graphs with broad, correlated degree distributions [16]. In this case, for a network of size *N*, the spectrum consists of a large number of eigenvalues (of order *N*) which lie within a disk in the complex plane, as well as a small number of outliers. One of these outliers lies along the real axis and moves to the right as the degree correlation increases. This result was found previously by one of our authors in studying the dynamics in networks of coupled oscillators [12]. It also agrees with and complements previous work on approximating the largest eigenvalue of random matrices [17]. Our work here suggests that this eigenvalue is related to the covariance term of the linearized operator in our mean-field model, Eq.(12), and is the relevant one for instabilities of the mean-field activity in networks of spiking neurons.

We showed that the degree covariances can be related to chain and reciprocal motifs in the network. For example, increasing the covariance between the excitatory in-degree and out-degree of E cells increases the number of *E* → *E* → *E* chains and reciprocally connected pairs of E cells. This is a particularly relevant example since the fraction of reciprocally connected pairs of E cells is a statistic which has been measured several times over the past few decades in cortical slice experiments [9–11]. It is robustly found that the fraction of reciprocal motifs is somewhere between 2 and 4 times what would be expected from an ER network. From our work here we can immediately conclude that one way to to achieve this is to allow for correlations between the excitatory in-degree and out-degree. Furthermore, the effect of this correlation on the mean-field is equivalent to increasing recurrent excitatory synaptic weights according to Eq. (6).

The mean-field model Eqs. (9) is heuristic. It is meant to describe the collective dynamics of large networks of E and I spiking neurons, but it is derived by assuming that a description of the dynamics in terms of mean firing rates (or synaptic outputs) is the relevant one. It may be that networks with broad, correlated degree distributions exhibit other modes of activity which cannot be captured in this framework, such as highly synchronous states. Indeed, it was shown in [13] that the out-degree of neurons strongly influences pairwise cross-correlations in synaptic inputs to neurons. It may be that this effect is enhanced if the out-degree is positively correlated to the in-degree. Also, other work studying the role of cortical motifs on dynamics in networks of integrate-and-fire networks using linear response theory has shown that both chain and divergence motifs can strongly alter pairwise correlations [18]. When the effects of synchrony are not significant for the population behavior, the mean-field model of Eqs. (9) can be used to study the effect of DDs on network dynamics with relatively little numerical or analytical effort.

## APPENDIX

### Degree Distributions

The degree of a given node in a graph is just the number of edges connected to it. In directed graphs such as neuronal networks, each node has two different degrees: an in-degree and an out-degree. The in-degree is the number of pre-synaptic inputs to a given neuron; the out-degree is the number of neurons which are post-synaptic to a given neuron. In this manuscript we consider networks in which the in-degree and out-degree of neurons are random variables drawn from a known joint-degree distribution.

#### Single population degree distribution

For the network consisting of a single population of *N* neurons, we denote the connectivity by the matrix W with entries *w_jk_* that are equal to one if there is a connection from neuron *k* onto neuron *j* and zero otherwise. We denote the in-degree of neuron *j* by 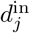 and its out-degree by 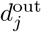. The degrees are defined in terms of the connectivity by 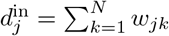 and 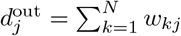.

To separate the influence of variations in the degree across the network from the effects of average connectivity, we normalize the degrees by the mean degree of the network, 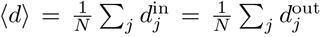. (The mean in-degree is identical to the mean out-degree because both sums are equal to the total number of connections on the network.) Throughout this manuscript, we will consider the normalized in-degree 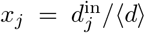 and out-degree 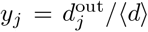, which we will refer to simply as degrees from now on.

For simplicity in the mathematical notation and analysis, we allow degrees to be continuous random variables and define the degree distribution ρ (*x, y*) as

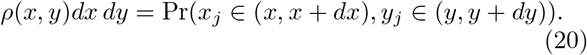

For the network consisting of a single population of neurons, we assume the network is statistically homogeneous in the sense that Eq. (20) holds independent of neuron index *j*.

#### Two population degree distribution

For the network consisting of *N_e_* excitatory neurons and *N_i_* inhibitory neurons, each neuron will have four degrees, as illustrated in Figure 10. Each neuron will have two in-degrees (an E in-degree and an I in-degree), as the in-degrees from each of the populations of neurons could differ. Similarly, each neuron will have two out-degrees (an E out-degree and an I out-degree), as the out-degrees to each population could differ.

**Fig. 10.**
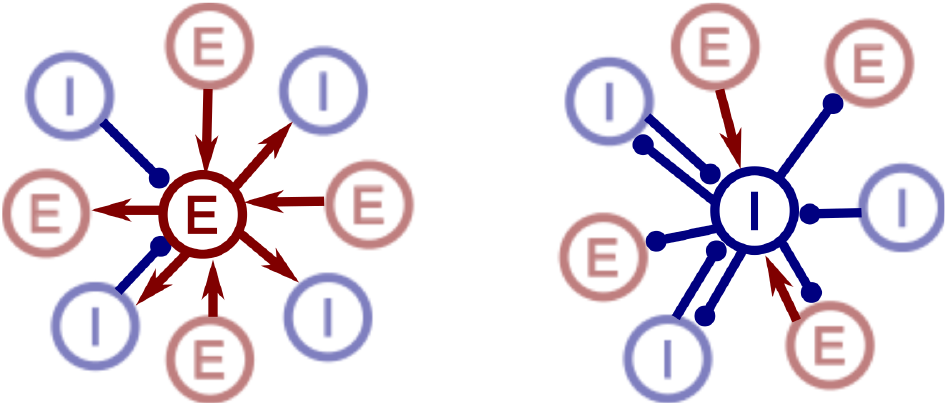
Illustration of degrees of excitatory and inhibitory neurons. To describe the degree of a single neuron requires four numbers: the number of connection to and from excitatory and inhibitory neurons. **Left**: The central excitatory neuron has three excitatory and two inhibitory incoming connections, giving it an E in-degree of 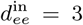 and an I in-degree of 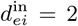. The neuron has one connection onto an excitatory neuron and three connections onto inhibitory connections, giving it an E out-degree of 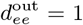 and an I out-degree of 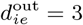. **Right**: The central inhibitory neuron has an E in-degree of 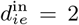, an I in-degree of 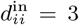, an E out-degree of 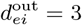, and an I out-degree of 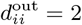.

For the two population case, we denote the components of the connectivity matrix *W* as 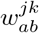, which equals one if neuron *k* from population *b* ∈ {*e, i*} synapses onto neuron *j* from population *a* ∈ {*e, i*}. Consider neuron *j* in population *a.* We denote its (unnormalized) in-degree from population *b* as

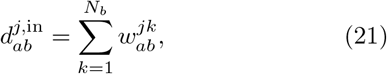

and its (unnormalized) out-degree onto population *b* as

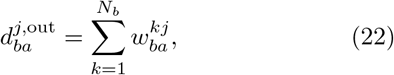

To factor out effects due to average connectivity, we will again use degrees which are normalized by the mean degree, defining

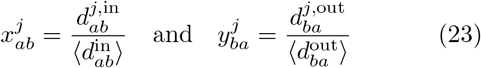

where 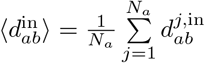 and 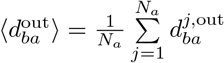.

As in the single population case, we’ll refer to the *x*’s and *y*’s simply as degrees and allow them to be continuous random variables. The four-dimensional degree distribution for neurons in population *a* ∈ {*e, i*} can be written as *ρa* (*x_ae_, x_ai_, y_ea_, y_ia_*). Denoting the two in-degrees as the vector **x**_*a*_ = (*x_ae_, x_ai_*) and the two out-degrees as **y**_*a*_ = (*y_ea_, y_ia_*), we can more compactly represent the degree distribution as *ρ_a_* (**x**_*a*_, **y**_*a*_).

### The firing rate model

To formulate equations governing the dynamics of the firing rate in the network that are based on the degree distribution of the neurons, we begin with the heuristic formalism developed by Wilson and Cowan [**?** ] where the total input to any given neuron is calculated by summing the activity of all neurons pre-synaptic to it. We reformulate the model in terms of the firing rate of neurons parameterized by their degree. This firing rate model will form the basis the mean-field models that are the central focus of this paper.

#### Single population firing rate model

For the single population model, if we let *r_j_* (*t*) be the firing rate of neuron *j* and *r ˙_j_* (*t*) denote its derivative, then one form of the Wilson-Cowan equations for the evolution of the firing rates is

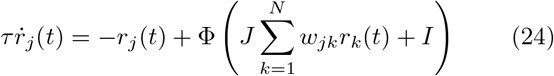

where *J* is a synaptic weight, *I* is the external current, and Φ is a sigmoidal nonlinearity giving the steady state firing rate of a neuron as a function of its input. The firing rate relaxes to its steady state value with a characteristic time constant τ.

In our approach, rather keeping track of the firing rate of neurons by their index *j*, we keep track of firing rates of neurons based on their in-degree *x* and out-degree *y* (we can say by their degree pair (*x, y*)). If we let *r*(*x, y, t*) be the firing rate of a neuron with degree pair (*x, y*), then to form an equation for *r* ˙ (*x, y, t*), we need only transform the sum over all neuron indices to an integral over all degree pairs (*x ˜*, *y ˜*). In this sum, we must weight the firing rate of each degree pair (*x ˜*, *y ˜*) by the probability density *ρ* (*x ˜*, *y ˜*) of that degree pair being present in the network. This self-averaging argument is valid if the network is large compared to the largest degree.

The remaining task is transforming the connectivity *w_jk_* into a representation in terms of degrees. We ignore the detailed structure of the network and simply assume that, given the degree distribution, all connections are made randomly and with equal probability. In this case, the probability of a connection from a neuron with degree pair (*x˜*, *y˜*) onto a neuron with degree pair (*x, y*) is proportional to the in-degree *x* of the post-synaptic neuron and the out-degree *y˜* of the presynaptic neuron. This result, illustrated in Fig. 11, can be seen intuitively since the likelihood of making a connection increases both with the number of out-going connections from the pre-synaptic neuron and the number of in-coming connections to the post-synaptic neuron. This form for the probability becomes exact when the degrees are small compared to the system size *N*. We absorb the proportionality constant into the coupling strength factor *J* and let the coupling strength from a neuron with degree pair (*x˜*, *y˜*) onto a neuron with degree pair (*x,y*) be *Jxy˜*.

**Fig. 11.**
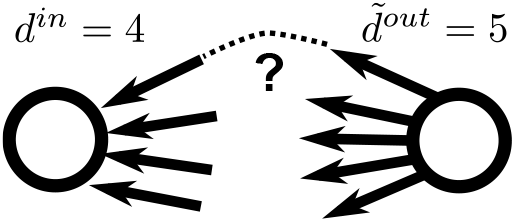
The probability of a connection is determined by the out-degree of the pre-synaptic and the in-degree of the post-synaptic. In this illustration, the pre-synaptic out-degree is *d*^out^ = 5 and the post-synaptic in-degree is *d*^in^ = 4, as represented by the arrows. The probability of a connection from the presynaptic neuron to the postsynaptic neuron is the probability that one of the right arrows is the same as one of left arrows, which is proportional to the product *d*^in^ *d̃*^out^.

With these conventions, we transform Eq. (24) into a self-consistent equation for the firing rate *r*(*x, y, t*) of a neuron with degree pair (*x,y*):

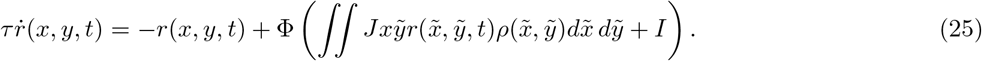

The recurrent input (the integral term) is just the firing rate of each neuron in the network, weighted by the probability of connection, and averaged over the degree distribution. Note that this input does not depend on the out-degree of the post-synaptic neuron, and so we can write

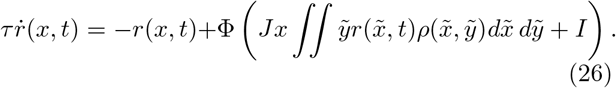

The effect of the out-degree, however, does not drop out of the equation in general. Specifically, pre-synaptic firing rates are weighted by the out-degree of the corresponding neurons. As we will see, this weighting can play an important role in the dynamics.

#### Two population firing rate model

The firing rate model for the EI network is analogous to the single population model, with two important differences. The first difference is that we now have two sets of equations, one for the firing rates *r_e_*(**x**_*e*_, *t*) of the excitatory neurons and one for the firing rates *r_i_*(**x**_*i*_, *t*) of the inhibitory neurons. As for the single population case, these firing rates just depend on the in-degrees, though for two population, each neuron has two different in-degrees. The second difference is that each firing rate equation has two input terms, one for the excitatory input and one for inhibitory input.

We let *J_ab_* be the coupling strength factor for connections from population *b* onto population *a*. Given that the probability of connection from a neuron *j* to a neuron *i* is proportional to the relevant out-degree of *j* and the relevant in-degree of *i*, the self-consistent firing rate equations for the EI network are

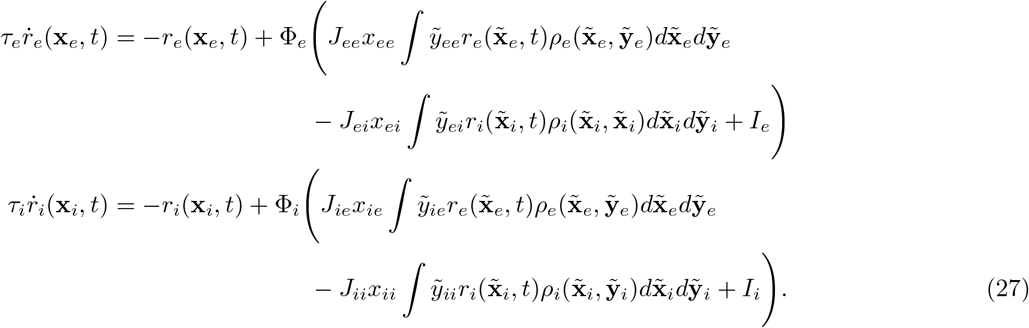

Since each neuron has both an E and an I in-degree, **x**_*a*_ = (*x_ae_, x_ai_*), and both an E and an I out-degree, **y**_*a*_ = (*y_ea_, y_ia_*), the integrals which determine the recurrent input in Eqs. (27) are actually quadruple integrals and *dx_a_ dy_a_* = *dx_ae_dx_ai_dy_ea_dy_ia_*.

### The integrate-and-fire network model

We implemented an integrate-and-fire network model using the Brian simulator [19]. The network consisted of 4000 excitatory neurons and 1000 inhibitory neurons with no excitatory-excitatory connections. The subthreshold dynamics of the voltage *V_e_^j^* (*V_i_^j^*) of neuron *j* of the excitatory (inhibitory) population were governed by the equations

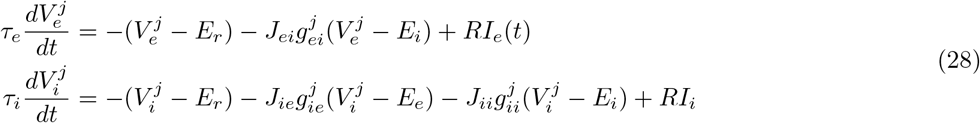

which were driven by normalized conductances that, in the absence of input, decayed exponentially according the equations 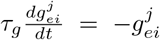, 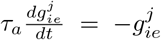 and 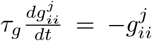. When the voltage of a neuron reached the threshold *υ*_th_, it was considered to have fired a spike. Its voltage was reset to *υ*_reset_ and held there for a refractory period of length *τ*_ref_; at the same time, conductances of its postsynaptic neurons were incremented by 1. Note that the network model did not include recurrent excitatory connections.

### Network structure

To generate EI-networks with correlated degree distributions, we used an extension of second order networks (SONETs) [12] to two populations. The SONET model is a random network model in which one can prescribe second order statistics (i.e., correlations) among certain edges in the network. These correlations can be viewed as specifying the frequencies of certain second order motifs (patterns of two edges in the network). If one fixes the average connectivity of the network, such motif frequencies correspond directly to covariances in the degree distribution [12]. Hence, we used the SONET model to generate networks with particular covariances in the degree distribution.

The SONET model is a probability distribution for an adjacency matrix *W* with components 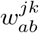 denoting connections from node *k* of population *b* to node *j* of population *b*. The SONET model prescribes the following statistics on the random components of *W*

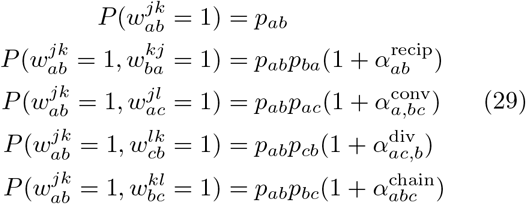

where (*j, a*), (*k, b*), and (*l, c*) could be any combination of distinct (node, population) combinations. Since the network has only two populations, some of the populations *a, b, c* ∈ {*e, i*} must be identical. Eq. (29) defines the 27 parameters of the two population SONET model: *p_ab_*, 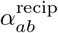, 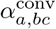, 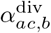, and 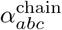. (There are only 27 parameters as 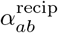, 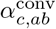 and 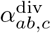 are symmetric in *a* and *b*.)

An illustration of the first and second order motifs of the SONET model is shown in Fig. 12. The code used to generate SONET networks can be found at github.com/dqnykamp/sonets.

**Fig. 12.**
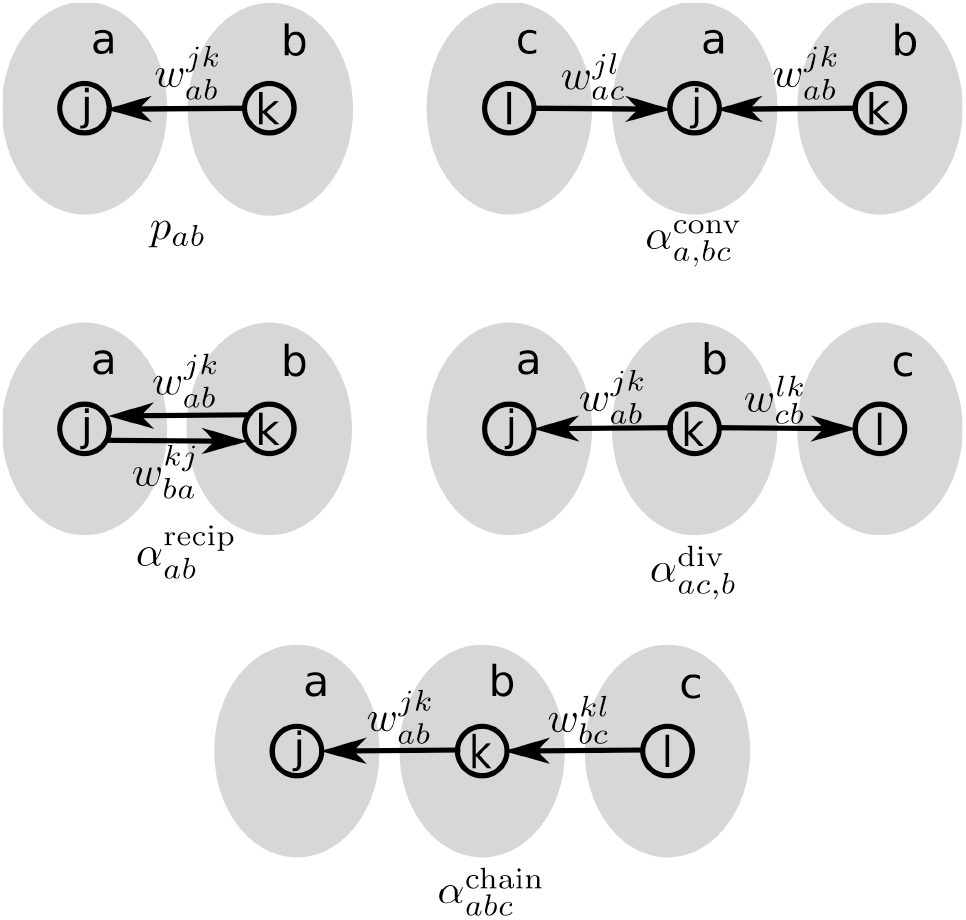
Illustration of the motifs underlying the SONET network model. The model contains one type of first order motif (a single edge represented by the parameter *p_ab_*) and four types of second order motifs: reciprocal (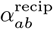), convergent (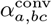), divergent (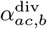), and chain (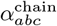) motifs. The relationship between the motif probabilities and the parameters is given in Eq. (29). Gray shaded regions represent populations *a, b,* and *c*, some of which must indicate the same population, as in this model, we have two populations: *a, b, c* ∈ {*e, i*}.

The SONET parameters 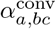, 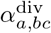, and 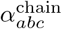 determine the degree distribution, as shown in [12] for a single population. Assuming large populations and large variance in-degrees, the convergent motif frequencies correspond to the covariances of the (normalized) in-degrees(*x_ae_,x_ai_*),

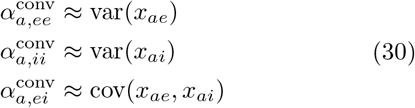

the divergent motif frequencies correspond to the covariances of the (normalized) out-degrees (*y_ea_, y_ia_*),

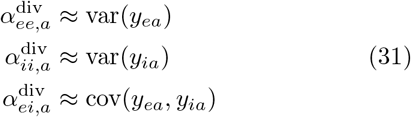

and the chain motif frequencies correspond to the covariances between the (normalized) in- and out-degrees,

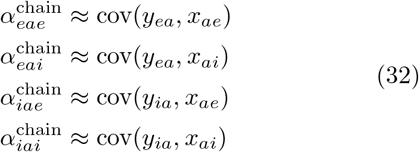

for *a* ∈{*e,i*}

## ACKNOWLEDGMENTS

A.R. acknowledges a Ramón y Cajal grant from the Spanish government and a project grant from the Spanish ministry of Economics and Competitiveness BFU2012-33413. A.R. has been partially funded by the CERCA progam of the Generalitat de Catalunya. Research also supported by National Science Foundation grant DMS-0847749 (D.Q.N, M.S, M.V.), the MBI Undergraduate Summer Research Program (D.F.), the Generalitat de Catalunya PIV-DGR 2010 program (D.Q.N). This work was funded by the Ministry of Economy and Competitiveness of Spain and the European Regional Development Fund (Ref: BFU2009-09537).

## References

[1] D. J. Amit and N. Brunel, Cerebral Cortex 7, 237 (1997).

[2] A. Roxin, N. Brunel, D. Hansel, G. Mongillo, and C. van Vreeswijk, J. Neurosci. 31, 16217 (2011).

[3] D. J. Amit and N. Brunel, Network 8, 373 (1997).

[4] G. Deco and E. Hugues, PLoS Comp. Biol. (2012).

[5] A. Litwin-Kumar and B. Doiron, Nat. Neurosci. 15, 1498 (202).

[6] R. Ben-Yishai, R. Lev Bar-Or, and H. Sompolinsky, Proc. Natl. Acad. Sci. USA 92, 3844 (1995).

[7] A. Compte, N. Brunel, P. S. Goldman-Rakic, and X.-J. Wang, Cerebral Cortex 10, 910 (2000).

[8] A. Roxin, N. Brunel, and D. Hansel, Phys. Rev. Lett. 94, 238103 (2005).

[9] H. Markram, Cereb. Cortex 7, 523 (1997).

[10] S. Song, P. J. Sjöström, M. Reigl, S. Nelson, and D. B. Chklovski, PLoS Biology 3, 507 (2005).

[11] R. Perin, T. K. Berger, and H. Markram, PNAS 108, 5419 (2011).

[12] L. Zhao, B. Beverlin, T. Netoff, and D. Q. Nykamp, Front. Comput. Neurosci. 5, 28 (2011).

[13] A. Roxin, Front Comput Neurosci 58, 8 (2011).

[14] Note that in [13] the neurons were parameterized by an “effective” in-degree *k* which can now be identified as the cumulative degree 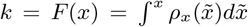. Therefore *dk* = *ρ_x_*(*x*)d*x* and the pre-factor *x* = *F*^−1^(*k*). This function of *k*, the inverse cumulative degree distribution, was called *J*(*k*) in [13], see e.g. Eq. 8 in that paper.

[15] P. Song and X. J. Wang, J. Neurosci. 25, 1002 (2005).

[16] J. Aljadeff, D. Renfrew, M. Vegué, and T. O. Sharpee, Phys. Rev. E 93, 022302 (2016).

[17] J. G. Restrepo, E. Ott, and B. R. Hunt, Phys. Rev. E 76, 056119 (2007).

[18] Y. Hu, J. Trousdale, K. Josic, and E. Shea-Brown, J. Stat. Mech. P03012, 1 (2013).

[19] D. F. Goodman and R. Brette, Front. Neurosci. 3, 192 (2009).

